# Evaluation of spontaneous seizure activity, sex-dependent differences, behavioral comorbidities, and alterations in CA1 neuron firing properties in a mouse model of Dravet Syndrome

**DOI:** 10.1101/2021.06.16.448684

**Authors:** Chelsea D. Pernici, Alexa Spink, E. Jill Dahle, Kristina J. Johnson, Cameron S. Metcalf, Peter J. West, Karen S. Wilcox

**Affiliations:** Epilepsy Therapy Screening Program (ETSP) Contract Site, University of Utah, Salt Lake City, Utah; Department of Pharmacology and Toxicology, University of Utah, Salt Lake City, Utah; Interdepartmental Neuroscience Program, University of Utah, Salt Lake City, Utah

## Abstract

Dravet syndrome (DS) is a rare childhood epilepsy disorder resulting in spontaneous, recurrent seizures (SRS) and behavioral co-morbidities. To facilitate the discovery and development of anti-seizure drugs for DS, the contract site of the NINDS Epilepsy Therapy Screening Program (ETSP) has continued to evaluate a mouse model of DS. *Scn1a*^*A1783V/WT*^ mice exhibited increased hyperactivity, thigmotaxis, and deficits in nest-building behavior. Ex-vivo brain slice electrophysiology experiments revealed increased excitability of hippocampal CA1 neurons specifically due to increased action potential firing frequency in response to brief depolarizations and decreased frequency of spontaneous GABAergic synaptic events. A video-EEG study revealed mice had on average, 1 seizure per day, with males seizing significantly more frequently than females. Increased proportion of seizure activity occurred during the dark phase of the light/dark cycle in both sexes. While clobazam, a drug commonly prescribed to patients with DS, had no effect on SRS activity at the tested doses, the seizure history and frequency observed in this study aids in determining the sample sizes and experimental timeline needed for adequately powered preclinical drug studies. Overall, this study provides a broad description of the *Scn1a*^*A1783V/WT*^ mouse and highlights the utility of this model in therapy discovery.

## Introduction

Dravet Syndrome (DS) is a devastating, rare, genetic epilepsy, with onset occurring in the first year of life (1). DS is a result of a mutation in the *SCN1A* gene, which encodes for the voltage-gated sodium channel, Nav_1.1_, ultimately causing loss-of-function(2, 3). Patients with DS typically present with a prolonged febrile seizure, followed by persistent, unprovoked, spontaneous seizures and cognitive decline. Symptoms are first apparent in the second or third year of life and cognitive decline continues to worsen throughout adolescence(4). DS is highly pharmacoresistant, and it is only recently that some treatments have become available(5). Previous work in murine models suggests that the Nav_1.1_ channel mutation affects inhibitory interneurons, causing them to be dysfunctional(6-9), leading to an electrical imbalance and results in run-away excitation and manifestation of a seizure. Discovering effective treatments is imperative for reducing the burdens associated with DS. To accomplish this, drug screening models with good construct, predictive, and face validity are needed. Historically, novel compounds have been screened using acute, evoked rodent seizure models, typically in naïve mice or rats. In addition to screening drugs against evoked seizures, it will be advantageous to screen drugs against spontaneous, recurrent seizures(10). Thus, preclinical drug screening models that better recapitulate aspects of epilepsy, such as spontaneous, recurrent seizures, are needed. Our previous work has shown a mouse model of DS with a knock-in *Scn1a* mutation has a 50% survival rate and hyperthermia-induced seizures. The hyperthermia-induced seizures are pharmacoresistant to a battery of prototype anti-seizure drugs, with the exception of clobazam, tiagabine, levetiracetam or when treated with a commonly used drug combination of clobazam, valproic acid, and stiripentol(11). However, it is unknown if spontaneous seizures demonstrate the same pharmacoresistance as evoked seizures.

An important goal at the contract site for the NINDS Epilepsy Therapy Screening Program (ETSP) is to have translatable models of therapeutic treatment effects in patients. Models which best recapitulate clinical observations are expected to offer improved translation. In DS, while patients experience febrile seizures, they are primarily affected by high spontaneous seizure burden and behavioral comorbidities. Therefore, we extend our knowledge of the *Scn1a*^A1783V*/WT*^ mouse model of DS by assessing behavioral and spontaneous seizure phenotypes. The goal of the present study was to better understand if this model recapitulates phenotypes seen in a clinical setting and establish an effective drug screening paradigm for spontaneous seizures. Mice with the *Scn1a* mutation had increased thigmotaxis and hyperactivity as compared to wild-type littermates. Brain slice electrophysiology experiments revealed increased excitability of CA1 neurons in the hippocampus and decreased frequency of spontaneous GABAergic synaptic events. We determined the frequency of spontaneous seizures, investigated the effect of an etiological relevant priming event to increase seizure burden and determined the time of day when seizure frequency was highest. Additionally, we sub-chronically administered clobazam, a first-line treatment for DS(1), to determine its efficacy against spontaneous seizures. Finally, the interictal spike analysis performed here sets the stage for future work utilizing closed loop strategies in neuromodulation of seizures. Overall, this study serves to optimize and determine the duration of treatment time required for future drug screening experiments.

## Results

### Scn1a^A1783V/WT^ mice have impaired nest building behavior, anxiety-like behavior, and thigmotaxis

Nest building behavior was evaluated beginning at post-natal day (PND) 28 in both male and female *Scn1a*^*A1783V/WT*^ mice and wild type littermates at 2 and 6 hr, and then every 24 hours for 7 days. Mice were singly housed for ease of nest building evaluation. At each time-point, *Scn1a*^*A1783V/WT*^ mice had significantly impaired nest building behavior as compared to the wild type littermates (**Figure 1A**). No differences were present between male and female *Scn1a*^*A1783V/WT*^ mice (**Figure S1)**.

**Figure 1.**
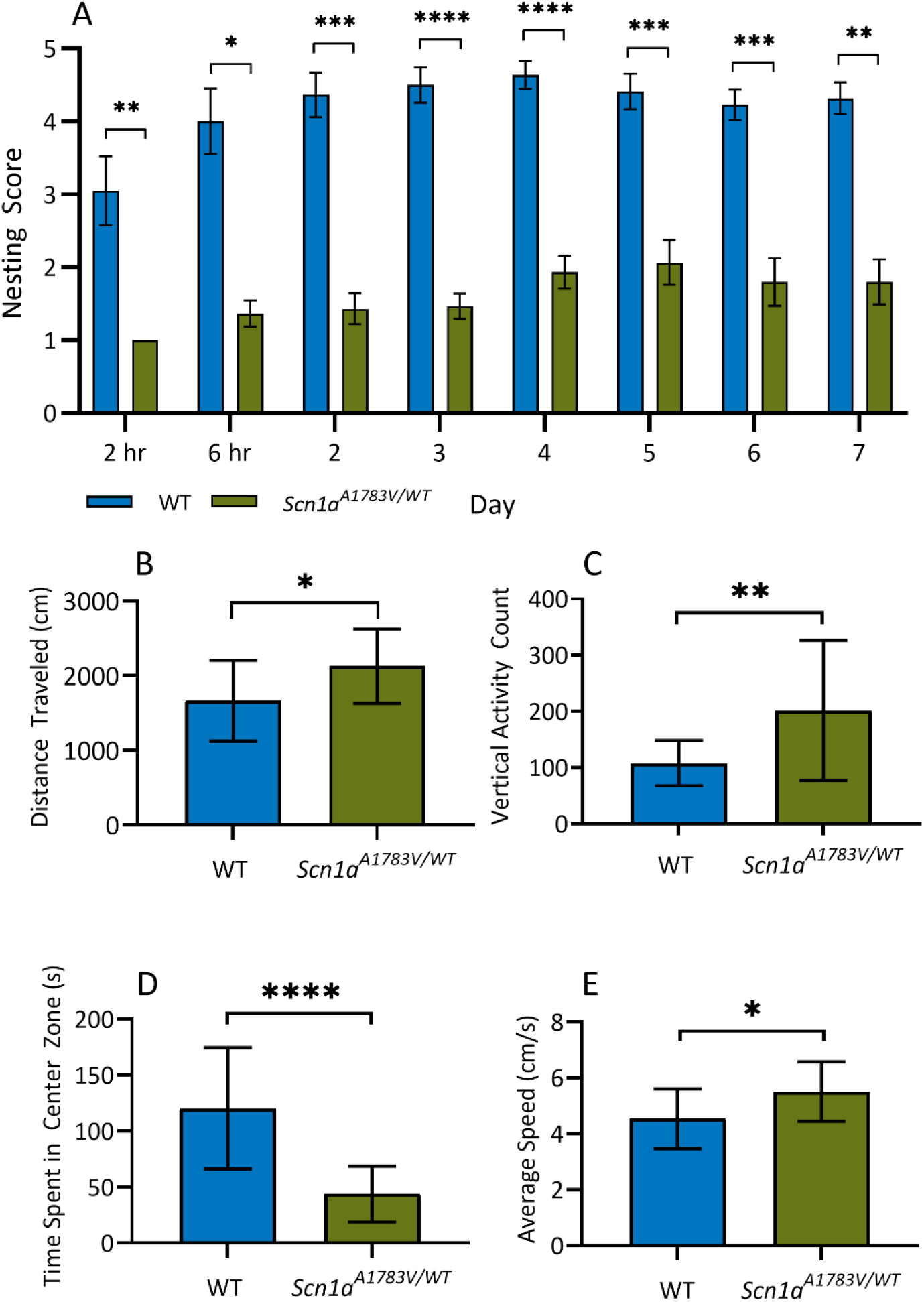
A) *Scn1a*^*A1783V/WT*^ (n=15) mice had significant nest building impairments as compared to wild type littermates (n=12) at each time-point (2 hr: p = 0.0049, 6 hr: p = 0.0114, Day 2: 0.002, Day 3: p<0.0001, Day 4: p< 0.0001, Day 5 p = 0.0009, Day 6: p = 0.0009, Day 7: p = 0.0012; Mann-Whitney, * p < 0.05, ** p < 0.01, *** p < 0.001, **** p < 0.0001 vs. WT) B) *Scn1a*^*A1783V/WT*^ mice traveled significantly farther (WT=1665 ± 540.8 cm, *Scn1a*^*A1783V/WT*^=2127 ± 133.6 cm, p=0.0224) and B) had a significantly higher average speed (WT= 4.54 ± 1.1 cm/s; *Scn1a*^*A1783V/WT*^ = 5.51 ± 1.1 cm/s, p=0.0196) than their wild type littermates. C) *Scn1a*^*A1783V/WT*^ spent signifcantly less time in the center zone (WT = 120.2 ± 54.2 s vs. *Scn1a*^*A1783V/WT*^ = 43.91 ± 24.9 s; p <0.0001) and D) had a significantly higher vertical activity count than their wild-type littermates (WT = 107.6 ± 40.3 vs. *Scn1a*^*A1783V/WT*^ = 201.5 ± 124.5 ; p=0.008). * p < 0.05, ** p <0.01, **** p < 0.0001; unpaired t-test; Data is presented as mean ± SD.

The open-field paradigm was used to evaluate several behavioral phenotypes present in both male and female P28 *Scn1a*^*A1783V/WT*^ mice. During a 10-minute evaluation period in an open-field arena, *Scn1a*^*A1783V/WT*^ mice traveled significantly farther and faster than their wild-type litter mates, indicative of increased hyperactivity (**Figure 1B-C**). Additionally, *Scn1a*^*A1783V/WT*^ mice spent significantly less time in the center of the open field, as well as had significantly higher vertical activity counts than their wild-type littermates, suggesting increased anxiety-like behaviors (**Figure 1D-E**). No differences were present between male and female *Scn1a*^*A1783V/WT*^ mice in the open field test (**Figure S2**). Thus, these experiments suggest that the *Scn1a*^*A1783V/WT*^ mice exhibit behavioral deficits consistent with those seen in both patients with DS(12-14) and other rodent and zebrafish models of DS(15-19).

### CA1 neurons in Scn1a^A1783V/WT^ mice have increased neuronal excitability and receive decreased sIPSC activity

Using the whole cell patch clamp technique, we determined if CA1 pyramidal neurons have altered excitability, dependent on age. Cells in the CA1 region of the dorsal hippocampus were current clamped in the whole-cell configuration and a graded series of hyperpolarizing and depolarizing current steps were used to examine electroresponsive membrane properties (**Figure 2 A-F**). There was no significant difference in membrane resistance (**Figure 2H**), membrane time constants (**Figure 2H**), resting membrane potential, or action potential dynamics (**Table 1**) between Scn*1a*^*A1783V/WT*^ and wild-type (*Scn1a*^*WT/WT*^) mice, independent of age (**Table S3**). However, there was a significant decrease in the inter-event interval between the first two action potentials resulting from a 200 pA current step for PND 18-25 *Scn1a*^*A1783V/WT*^ as compared to wild-type littermates. In PND >30 *Scn1a*^*A1783V/WT*^ mice, there was a significant decrease in the inter-event interval resulting from a 50 pA current step and non-significant trend of decreasing inter-event intervals for the other current steps (**Figure 2I, Table S2**). Given that there was some indication of increased intrinsic excitability in CA1 neurons of *Scn1a*^*A1783V/WT*^ mice, changes in excitatory or inhibitory input to the CA1 region were additionally investigated (**Figure 3A-B, E-F**). Recordings of sEPSCs showed that neither amplitude nor frequency of events were significantly altered in PND 30-60 *Scn1a*^*A1783V/WT*^ mice (**Figure C-D**). However, the relative frequency of sIPSC events in PND 30-60 *Scn1a*^*A1783V/WT*^ was significantly reduced as compared to wild types, while amplitude was unaffected (**Figure 3G-H**). This is consistent with prior work in other animal models of DS, where decreased sodium channel expression in interneurons results in a concomitant decrease in sIPSC frequency(20).

**Figure 2.**
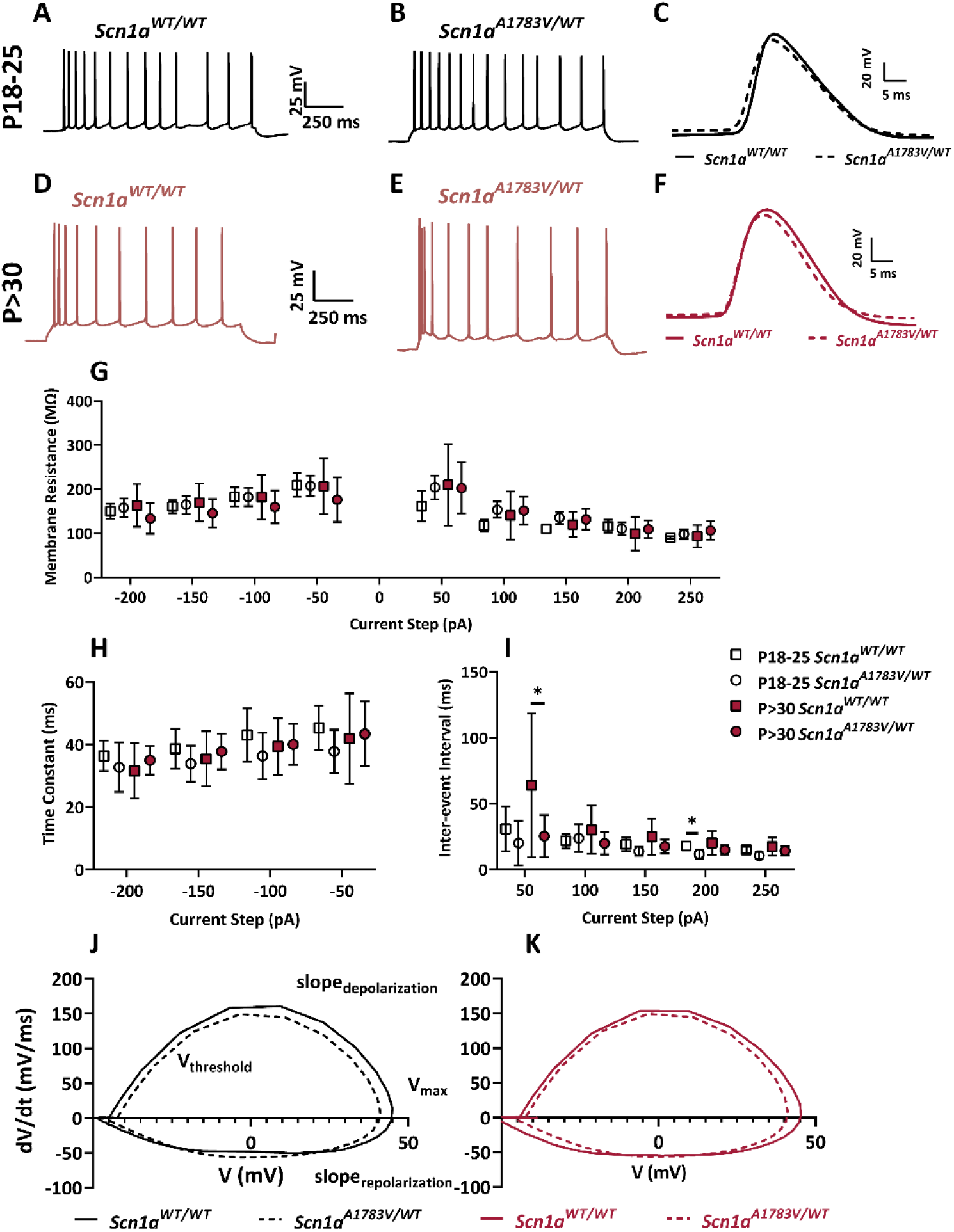
A-B) Representative traces of voltage responses to a 50 pA current step for PND 18-25 *Scn1a*^*WT/WT*^ and *Scn1a*^*A1783V/WT*^ mice and D-E) PND >30 *Scn1a*^*WT/WT*^ and *Scn1a*^*A1783V/WT*^ mice. C) Representative trace of a single action potential for PND 18-25 *Scn1a*^*WT/WT*^ and *Scn1a*^*A1783V/WT*^ mice and F) PND >30 *Scn1a*^*WT/WT*^ and *Scn1a*^*A1783V/WT*^ mice. G) There were no significant differences in measured membrane resistance between *Scn1a*^*WT/WT*^ and *Scn1a*^*A1783V/WT*^ mice, independent of age or current-step induced changes in voltage. H) There were no significant differences in time constants between *Scn1a*^*WT/WT*^ and *Scn1a*^*A1783V/WT*^ mice, independent of age or current-step induced changes in voltage. I) Inter-event interval was measured as the time between the first two action potential peaks at each current step. PND 18-25 *Scn1a*^*1783V/WT*^ mice had significantly shorter inter-burst intervals at 100 pA as compared to wild type littermates (p=0.0283). PND >30 *Scn1a*^*A1783V/WT*^ mice had significantly shorter inter-burst intervals at the 50 pA current steps (p = 0.0365, unpaired t-test) J) Representative phase plots for PND 18-25 and PND >30 *Scn1a*^*WT/WT*^ and *Scn1a*^*A1783V/WT*^ mice. (PND 18-25 *Scn1a*^*WT/WT*^ N=2 mice/4 cells, *Scn1a*^*A1783V/WT*^ N=3 mice/5 cells; PND >30 Scn1a^WT/WT^ N=4 mice, 13 cells, *Scn1a*^*A1783V/WT*^ N=4 mice/13 cells; unpaired t-test; individual p-values are listed in Table S1-S3)

**Table 1.** Action potential properties of CA1 excitatory neurons in *Scn1a*^*WT/WT*^ and *Scn1a*^*A1783V/WT*^ mice.

|  | P18-25 |  |  | P>30 |  |  |
| --- | --- | --- | --- | --- | --- | --- |
|  | *Scn1a*<sup>WT/WT</sup><br>(N=2 mice/4<br>cells) | *Scn1a*<sup>A1783V/WT</sup><br>(N=3 mice, 5<br>cells) | p-value | *Scn1a*<sup>WT/WT</sup><br>(N=4 mice/13<br>cells) | *Scn1a*<sup>A1783V/WT</sup><br>(N=4 mice, 13<br>cells) | p-value |
| **Resting Potential**<br>**(mV)** | -55.4 ± 4.5, | -58.8 ± 2.0 | 0.1686 | -58.0 ± 4.8 | -57.4 ± 3.0 | 0.7164 |
| **Threshold (mV)** | -43.6 ± 1.9 | -43.3 ± 4.2 | 0.9116 | -44.1 ± 3.0 | -44.1 ± 3.0 | 0.9901 |
| **Max Depolarization**<br>**Slope (mV/ms)** | 164.4 ± 40. | 158.4 ± 42.7 | 0.8350 | 171.3 ± 50.5 | 164.0 ± 39.2 | 0.6873 |
| **Max Depolarization**<br>**(mV)** | 44.9 ± 9.2 | 41.3 ± 10.3 | 0.6062 | 47.9 ± 13.6 | 45.5 ± 10.1 | 0.8867 |
| **Max Repolarization**<br>**Slope (mV/ms)** | -51.1 ± 8.0 | -48.1 ± 10.7 | 0.6517 | -52.6 ± 10.7 | -52.1 ± 7.4 | 0.6823 |

*Data represented as mean ± SD. Unpaired t-test*

**Figure 3.**
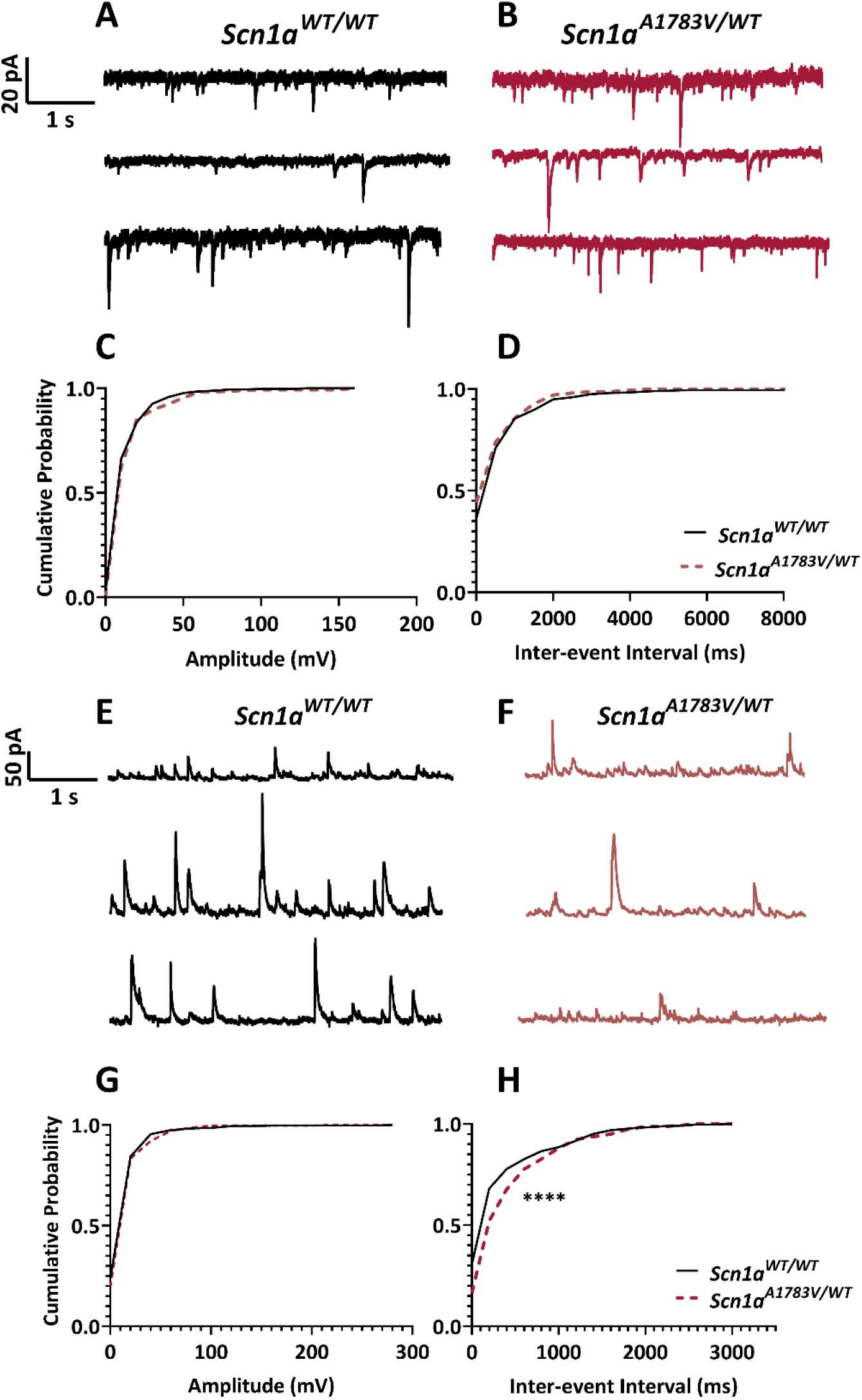
A-B) Representative traces of sEPSCs for P>30 *Scn1a*^*WT/WT*^ and *Scn1a*^*A1783V/WT*^ mice. There was no significant difference between the C) amplitude (p=0.1165) or D) frequency (p=0.0800) of sEPSC events (Kolmogorov-Smirnov test, *Scn1a*^*WT/WT*^ N=5 mice/7 cells, *Scn1a*^*A1783V/WT*^ N=5 mice/6 cells). E-F) Representative traces of sIPSC recordings for P>30 *Scn1a*^*WT/WT*^ and *Scn1a*^*A1783V/WT*^ mice. G) There was no significant difference in the amplitude of sIPSC events (p=0.0523) but there was a significant decrease in the frequency of events in the *Scn1a*^*A1783V/WT*^ mice as compared to wild-type (p<0.0001) (Kolmogorov-Smirnov test, Scn1a^WT/WT^ N=5 mice/7 cells, *Scn1a*^*A1783V/WT*^ N=5 mice/6 cells).

### Scn1a^A1783V/WT^ mice exhibit spontaneous seizure activity and males seize significantly more frequently than female mice

Both *Scn1a*^*A1783V/WT*^ and wild type littermates were implanted with a cortical, bipolar electrode and chronically monitored with video-EEG to determine if this model exhibited electrographic seizures and if the frequency of seizures was sufficient for drug studies. *Scn1a*^*A1783V/WT*^ had evidence of spontaneous seizure activity over a 14 day recording period, with the exception of two mice (**Figure 4A&B, Figure 5A-C**), which didn’t have a seizure until after an etiological priming event (**Figure 6)**. However, the other 96% of mice exhbited a seizure without a priming event. Seizures were defined as 2x the amplitude of baseline, lasting longer than 10 seconds followed by post-ictal depression (**Figure 4A&B)**. Wildtype littermates (n=10) had normal EEG activity and seizures were never observed. Daily seizure frequency was averaged over 14 days.

**Figure 4).**
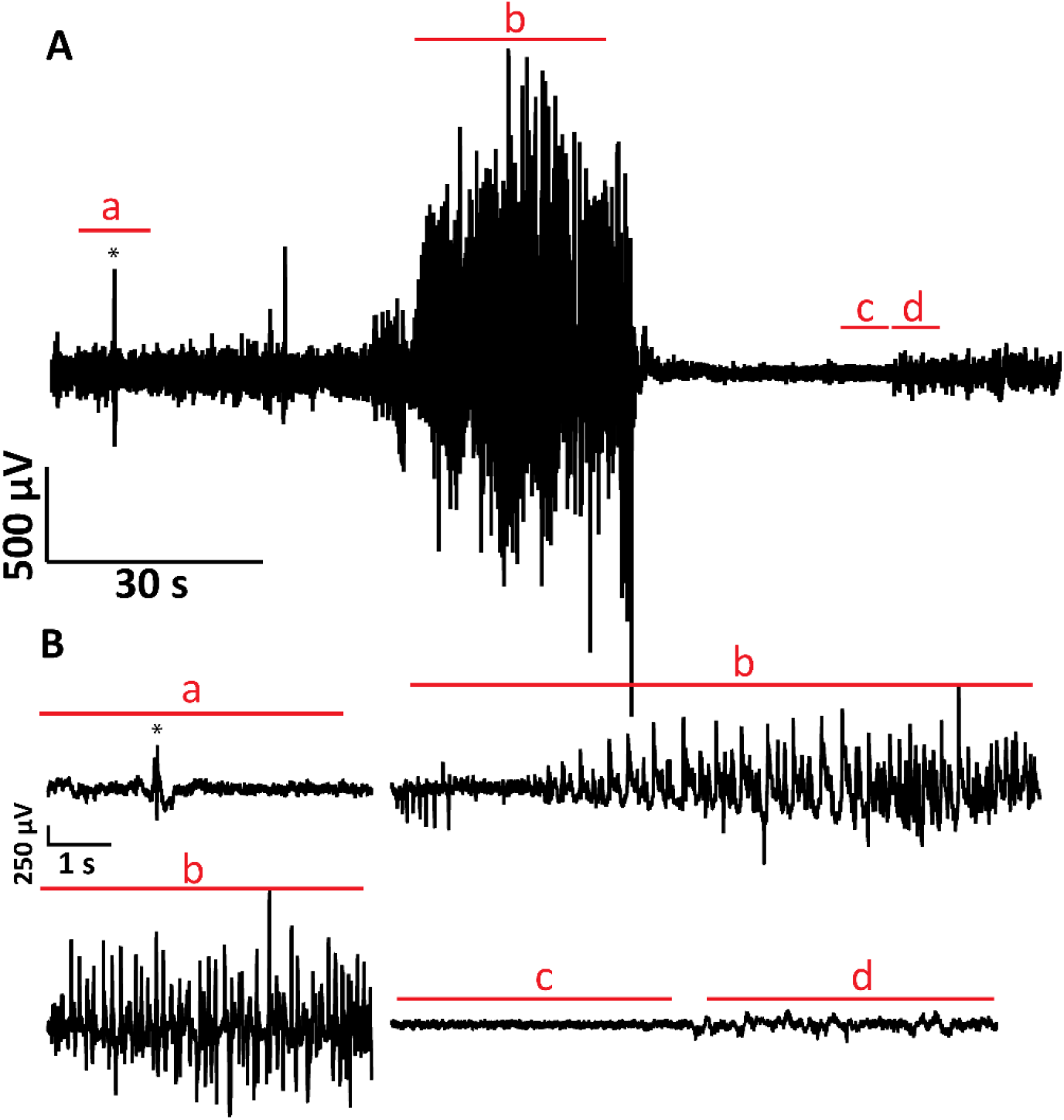
Mice were implanted with a cortical electrode and monitored with 24 hr, 7 day/week video-EEG (n=50). A) Representative trace of video-EEG recording from a *Scn1a*^*A1783V/WT*^ mouse. B) Expanded view of video-EEG trace. Lower case letters denote the following: a) baseline, normal activity with the * denoting an interictal spike. b) spontaneous seizure event with behavioral component c) post-ictal depression d) return to normal activity.

**Figure 5.**
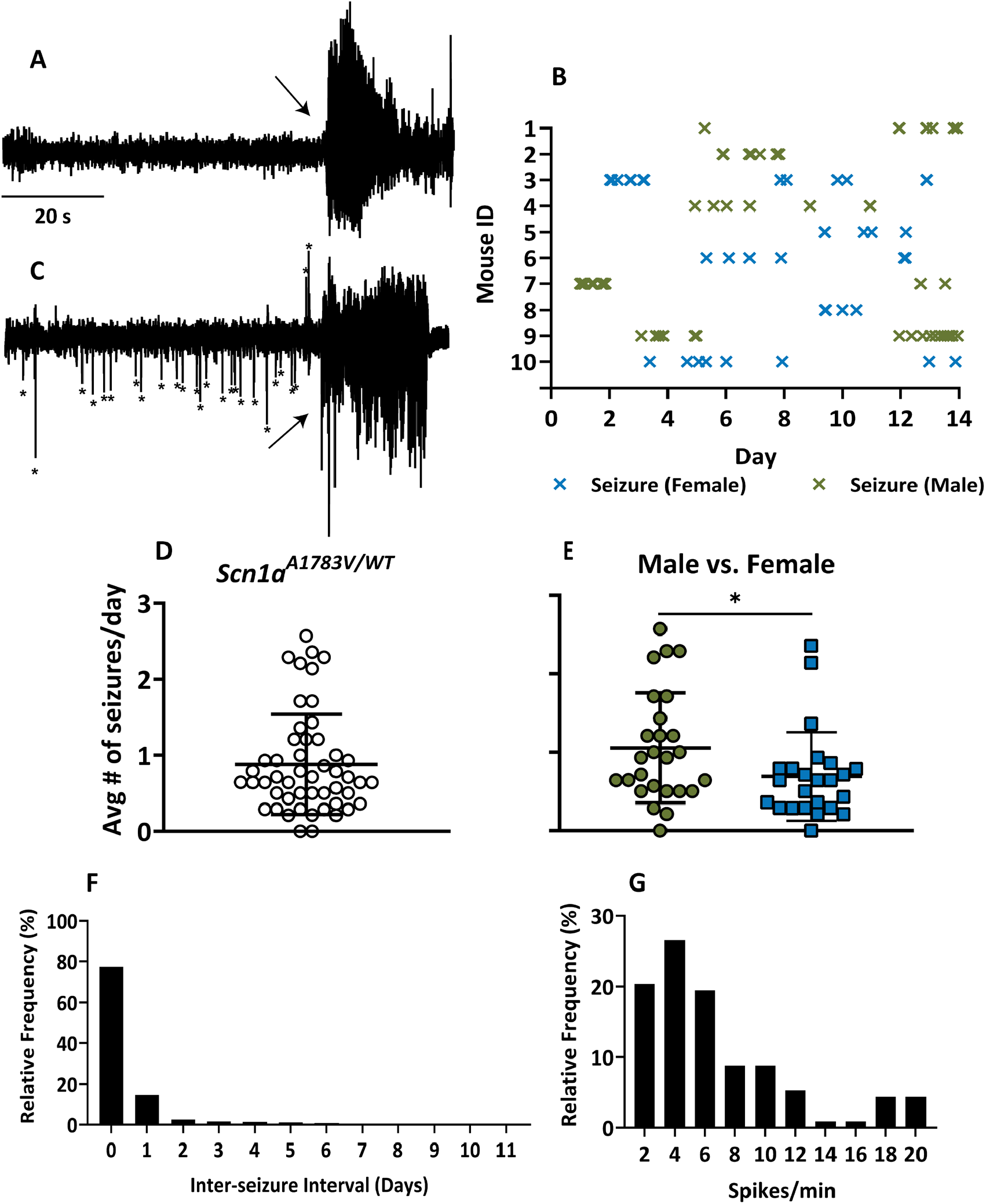
A) Representative trace of a spontaneous seizure. Seizures were classified as activity 2x the amplitude of baseline, lasting longer than 5 seconds, and displaying an increase in frequency. Arrow denotes beginning of ictal event. B) Representative plot of seizure events. Each ‘X’ denotes a spontaneous seizure. C) Representative trace of seizure preceded by an increase in inter-ictal events (* denotes inter-ictal events, arrow denotes beginning of ictal event) D) *Scn1a*^*A1783V/WT*^ mice have on average, 0.9 ± 0.7 spontaneous seizures per day, when measured over a 14-day period (n=50). E) Male *Scn1a*^*A1783V/WT*^ exhibit significantly increased seizure frequency (1.1 ± 0.7 seizures/day, n=26) as compared to female mice (0.7 ± 0.6 seizures/day, n=24) (p=0.0362, Mann-Whitney). F) Histogram of inter-seizure intervals (days) G) Approximately 76% of randomly selected seizures were preceded by a G) average of 6.7 ± 4.8 spikes/min.

**Figure 6.**
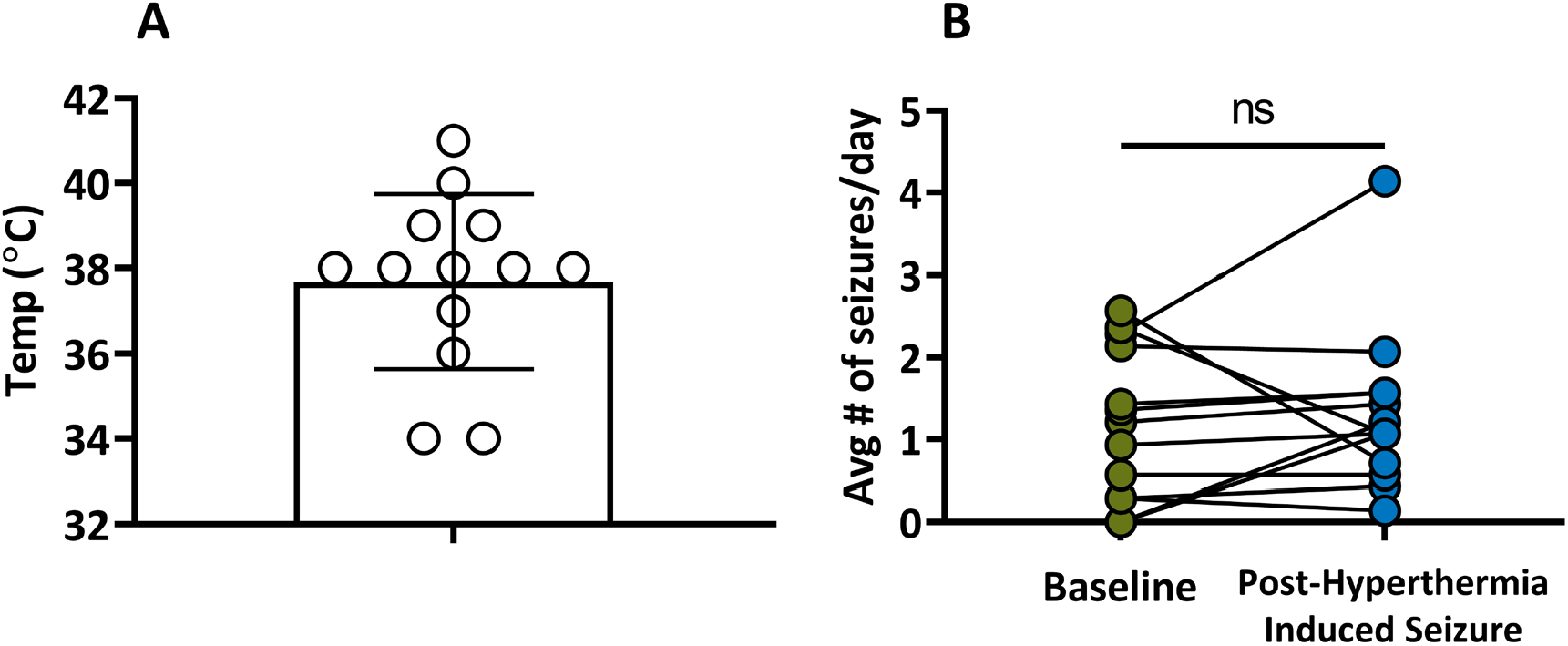
A) The average temperature at which mice had hyperthermia-induced seizures was 37.7 ± 2.1°C. Following the hyperthermia-induced seizure testing, video-EEG was collected for another 14 days. B) The addition of the hyperthermia-induced seizure did not significantly increase the daily seizure frequency as compared to baseline over a 14 day recording period (p = 0.4229, Wilcoxon matched-pairs; Baseline: 1.18 ± 0.9, Post Hyperthermia-induced seizure: 1.3 ± 0.9, n = 13)

*Scn1a*^*A1783V/WT*^ mice (n=50) had approximately 0.9 ± 0.7 seizures per day (**Figure 5D**). Synchronized video-EEG showed the behavioral component of the seizures included tonic-clonic seizures and “popcorn” behavior. All recorded seizures exhibited this behavioral component. While the average frequency was near 1 seizure/day, note that any mouse could go several days without seizing (**Figure 5B**). When data was evaluated based on sex, male *Scn1a*^*A1783V/WT*^ mice (1.1 ±0.7) had significantly more seizures per day than female *Scn1a*^*A1783V/WT*^ mice (0.7 ± 0.6) (**Figure 5E**). Interestingly, in a previous study, we noted that female mice had a greater mortality than male mice (68% vs 40%)(11). On average, each seizure lasted 34.7 ± 7.4 s (n=50), with no signifcant difference in seizure length between males (33.6 ± 6.4 s) and females (35.9 ± 8.2 s) (p=0.2619, unpaired t-test). Additionally, we calculated the inter-seizure event and found that 77.3% of the seizures occurred within 24 hrs of each other, with a smaller majority within 48 hr (14.6%), suggesting clustering activity (**Figure 5F**). We further investigated the clustering of seizures and found that 29/50 of the recorded mice experience at least one cluster during the 14-day recording period. A cluster was defined as one or more seizures per day for at least 3 days and at least 5 seizures in a cluster period(21). On average, each cluster included 10.3 ± 6.8 seizures. Mice had at least one seizure a day for 2.6 ± 1.4 consecutive days and had, on average, 4.2 ± 2.8 consecutive days without a seizure (**Table 2**). There were no significant differences in the clustering activity between male and female mice however, 70% of male mice had a seizure cluster, while only 45.4% of female mice had a seizure cluster in a 14-day period (**Table S4**). Finally, increases in inter-ictal spiking prior to seizure onset were observed during data analysis. A subset of seizures (n=150) were randomly selected and inter-ictal spikes in the 2 minute period immediately before an electrographic seizure were counted. This data set revealed that approximately 76% of seizures are preceded by inter-ictal spiking at a rate of 6.7 ± 4.8 spikes/min, while the other 24% of seizures had no evidence of inter-ictal spike activity (**Figure 5G)**.

**Table 2.** Cluster analysis for spontaneous recurrent seizures (SRS) in *Scn1a*^*A1783V/WT*^.

|  | **Consecutive**<br>**Days with SRS** | **Consecutive**<br>**Days without**<br>**SRS** | **# of mice with**<br>**clusters/# of**<br>**mice without**<br>**Clusters** | **SRS # per**<br>**Cluster** | **Cluster**<br>**Duration**<br>**(days)** |
| --- | --- | --- | --- | --- | --- |
| **Mean ± SD** | 2.3 ± 1.4 | 4.2 ± 2.8 | 29/50 | 10.3 ± 6.8 | 3.7 ± 0.9 |
| **Median** | 2.0 | 4.0 | ----- | 9.0 | 3.0 |
| **Max** | 6.0 | 14.0 | ----- | 35.0 | 6.0 |
| **Min** | 0.0 | 0.0 | ----- | 4.0 | 3.0 |

### Single hyperthermia-induced seizures do not increase spontaneous seizure frequency in Scn1a^A1783V/WT^ mice

Previous work has shown that a single hyperthermia-induced seizure increases spontaneous seizure frequency in a model of DS^40^. To determine if we could increase the average daily seizure frequency of *Scn1a*^*A1783V/WT*^ mice, a single hyperthermia-induced seizure was administered as a “priming” event. Baseline video-EEG data was collected for 14 days in N=13 mice and on Day 14, a seizure was induced by slowly raising the core temperature of the animal. Following this etiological priming event, video-EEG was collected and analyzed for an additional 14 days. All tested *Scn1a*^*A1783V/WT*^ mice seized at an average temperature of 37.7 ± 2.1 °C (**Figure 6A**). Both female and male mice were used as previous data has shown no difference in the temperature at which mice seize(11). A single hyperthermia-induced seizure event did not significantly increase the average daily seizure frequency in the cohort of mice tested over the two-week recording period (**Figure 6B**)

### Seizure frequency in Scn1a^A1783V/WT^ mice is dependent on time of day

The time of day at which mice seized was evaluated to determine if there was a day/night pattern associated with a proportion of seizure events. Seizures were first classified as occurring during either the light phase (between 6:00-18:00) or during the dark phase (18:00-6:00) of their light/dark cycle. A significantly higher proportion of seizures occurred during the dark phase as compared to the light phase (**Figure 7A**). The seizure events were further classified into 6-hr bins to determine if a higher proportion of seizures occurred during the first or second half of the dark phase. A significantly greater proportion of seizure events occurred between 18:00-24:00 hr than between 6:00-12:00 hr, 12:00-18:00 hr, or 0:00-6:00 hr (**Figure 7B**). There was no significant difference in seizure incidence between the light or dark phase between males and females (**Figure S3A**). When data was broken down into 6-hr bins, males seized significantly more between 18-24 hr than any other time. Additionally, during the 18-24 hr bin, males had a significantly higher proportion of seizures than females (**Figure S3B**).

**Figure 7.**
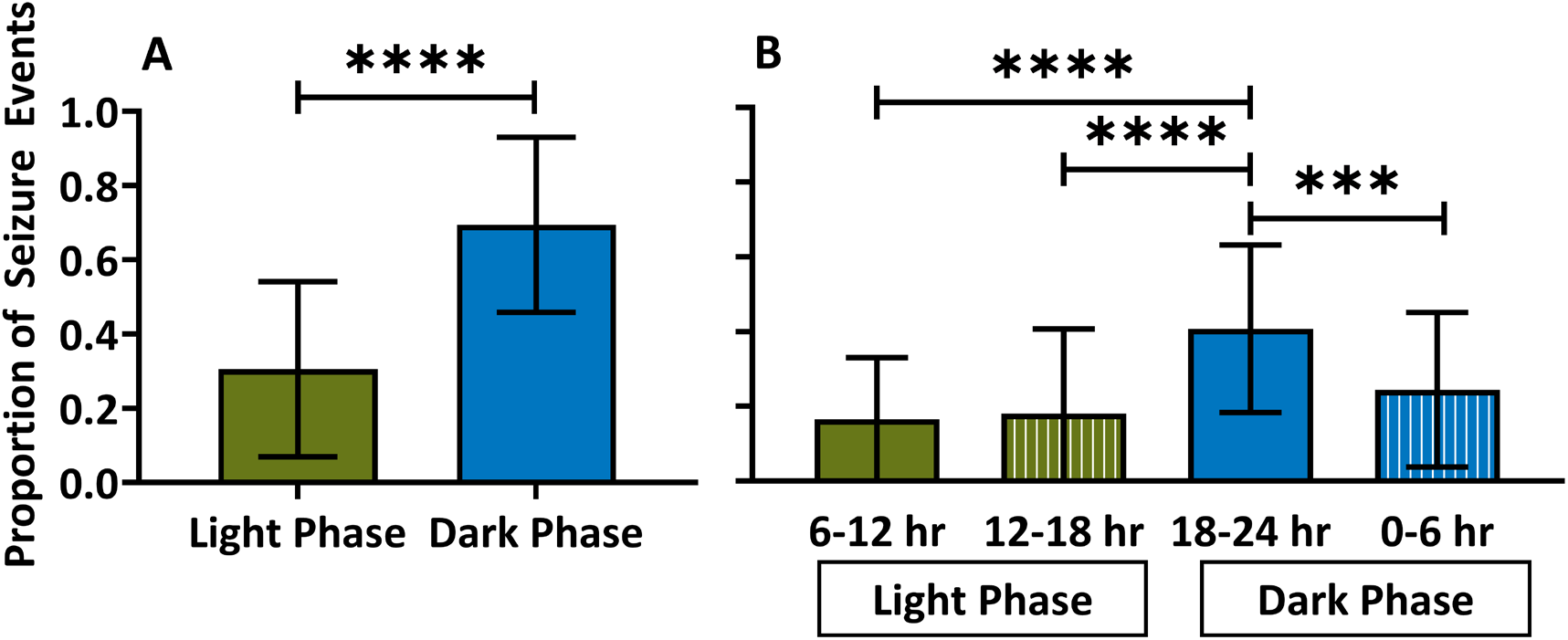
A) *Scn1a*^*A1783V/WT*^ mice had a significantly higher proportion of seizure events during the dark phase of their 12 hr light/12 hr dark cycle (Light: 0.33 ± 0.3; Dark: 0.67 ± 0.3; p < 0.0001) **** p < 0.0001, Mann-Whitney B) Seizure events were broken down into 6-hr bins and compared. *Scn1a*^*A1783V/WT*^ mice had a significantly higher proportion of seizure events between 18:00-24:00 than any other time-period (6-12 hr: 0.17 ± 0.2; 12-18 hr: 0.18 ± 0.2; 18-24 hr: 0.40 ± 0.2; 0-6hr: 0.25 ± 0.2, p = 0.0008 vs. 0-6 hr, p < 0.0001 vs. 6-12 hr, p < 0.0001 vs. 12-18 hr); *** p < 0.001, **** p < 0.0001; Ordinary one-way ANOVA

### Clobazam, at the tested doses, had no effect on seizure frequency

Clobazam is administered to patients with DS as a first-line treatment(1, 22). Thus, we tested CLB, at two different doses 5 mg/kg and 7.5 mg/kg. CLB was sub-chronically administered via i.p. injections, once a day for 10 days to determine if this compound significantly reduced seizure frequency in video-EEG recorded *Scn1a*^*A1783V/WT*^ mice. Mice in the study received both vehicle and drug in a cross-over scheme. Drug or vehicle was administered for 10 days, followed by a 7-day washout, and then groups were reversed for 10 days. At both a dose of 5 mg/kg and 7.5 mg/kg (i.p.), CLB did not significantly reduce seizure frequency as compared to vehicle treatment over a 10-day period (**Figure 8A&B**).

**Figure 8.**
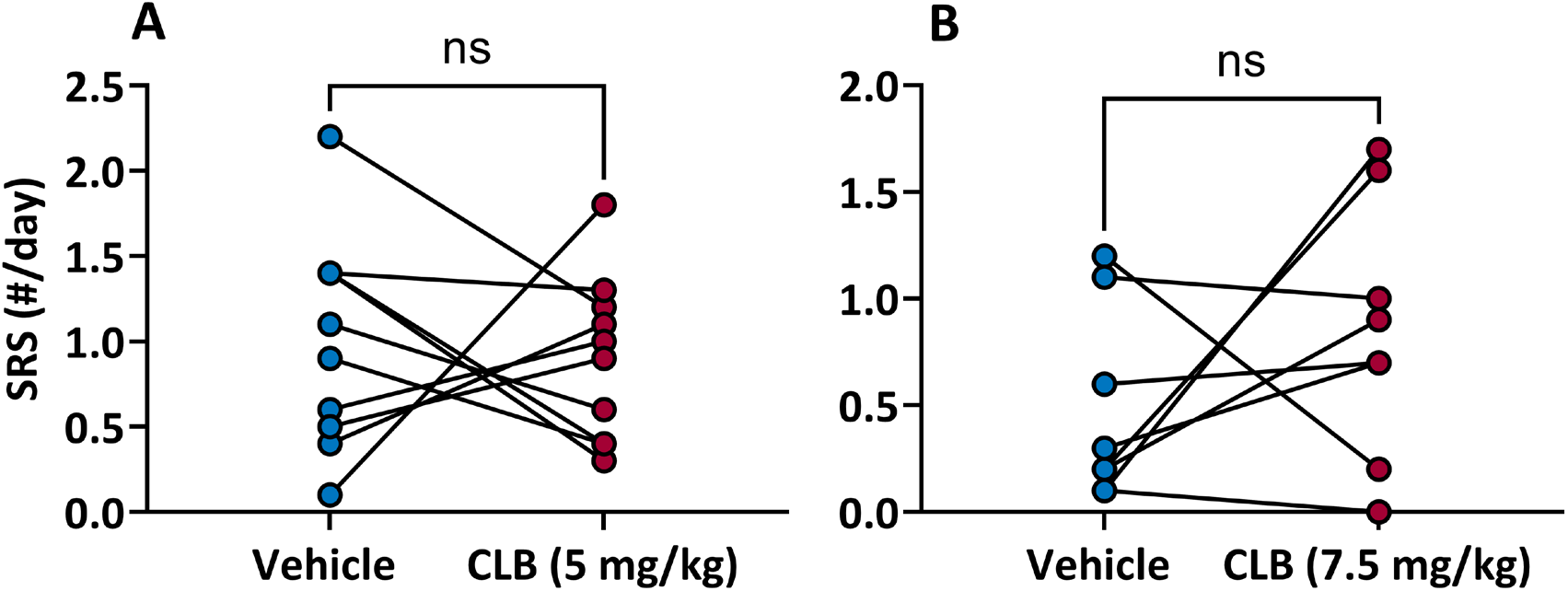
A). CLB (5 mg/kg, i.p.) did not significantly reduce seizure frequency when administered once a day over a 10-day period as compared to vehicle treatment (Vehicle: 1.00 ± 0.6; CLB (5mg/kg): 0.90 ± 0.5; p = 0.5352, Wilcoxon test) B) CLB (7.5 mg/kg) had no effect on seizure frequency when administered daily for 10 days as compared to vehicle treatment (Vehicle: 0.48 ± 0.4; CLB (7.5 mg/kg): 0.85 ± 0.6; p = 0.3125, Wilcoxon Test). Data presented as mean ± SD.

## Discussion

Our previous work demonstrated that the *Scn1a*^*A1783V/WT*^ mouse model of DS has a 50% survival rate and hyperthermia-induced seizures which are refractory to a battery of prototype anti-seizure drugs(11). The present study greatly extends our knowledge of the *Scn1a*^*A1783V/WT*^ mouse by systematically evaluating several attributes of this model, including behavioral comorbidities, spontaneous seizure activity, electrophysiological properties, and lack of sensitivity of spontaneous seizures to CLB. Of note, for the first time in this mouse model of DS, we have demonstrated sex dependent differences in seizure frequency, the occurrence of increased interictal spike frequency for two minutes prior to the seizures which will allow for predicting the occurrence of the next seizure, and the presence of seizure clusters. While the seizures did not respond to CLB at the doses used as hypothesized, the study provides a rational for future experiments, in terms of route of drug administration, drug dose timing, and length of pharmacological experiments.

In humans, the most reported behavioral comorbidities in patients with DS include hyperactivity, attention deficits, learning difficulties, and motor disorders(12-14). Similarly, mouse models of DS, including the *Scn1a*^*A1783V/WT*^ mouse model of DS, exhibit behavioral comorbidities. Evaluation in the open field showed *Scn1a*^*A1783V/WT*^ mice had significantly increased hyperactivity and anxiety-like behavior compared to wild type littermates and had poor nest building performance (**Figure 1**). This has also been shown by others investigating the *Scn1a*^*A1783V/WT*^ mice(17, 23) and in the *Scn1a*^*RX/+*^ and the *Scn1a*^*tm1Kea*^ mouse models(15, 16, 18). Poor nest building behavior is an early predictor of behavioral deficits and is an indicator of impaired activities of daily living(24). In addition to mouse models, Scn1ab mutant zebrafish also exhibit increased anxiety-like behavior in an open field(19). Models which recapitulate multiple aspects of the disorder are important to ensure face validity but also offers an opportunity to better understand if the comorbidities arise from the genetics, as an effect of multiple seizures, or as a result of ASDs.

DS is most often caused by a genetic mutation in the SCN1A gene, which encodes for the voltage gated sodium channel, Nav_1.1_. Evidence suggests that these mutations primarily affect inhibitory interneurons, leading to a loss of function in these cells, and causing an imbalance between excitation and inhibition(25). While our results show that most electroresponsive membrane properties were consistent between *Scn1a*^*A1783V/WT*^ and wild-type littermates, independent of age, *Scn1a*^*A1783V/WT*^ mice exhibited increased AP frequency during initial moments of depolarization (**Figure 2I**). Since all other electroresponsive membrane properties in these neurons were unaltered, the mechanism for this increase in burst-like AP frequency in CA1 neurons from DS mice remains to be determined. However, this increased AP frequency is consistent with what others have reported(7, 26). Furthermore, deficits in the Nav1.1 channel impair excitability of GABAergic interneurons(7, 9, 20, 25). We have shown this is consistent in the *Scn1a*^*A1783V/WT*^ mouse model as sIPSC frequency in area CA1 was decreased (**Figure 3H**). Thus, the *Scn1a*^*A1783V/WT*^ model of DS displays electrophysiological properties in area CA1 consistent with what is seen in other mouse models(20, 26-28).

A frequently reported limitation of mouse models of epilepsy, especially in terms of genetic models, is the low frequency of spontaneous seizure events. A goal of this study was to determine if the *Scn1a*^*A1783V/WT*^ mouse model exhibited spontaneous seizure activity and if the frequency of seizures was adequate for appropriately powered drug studies to evaluate novel compounds. We show that *Scn1a*^*A1783V/WT*^ mice have approximately 1 seizure per day, when measured over a 2-week time-period, and based on the inter-seizure interval, the seizures exhibit clustering properties (**Figure 4 and Table 2**). Male mice have significantly more seizures per day than females, yet our previous work has shown that females have increased mortality as compared to males (**Figure 5E**)(11). This was unexpected, as given the high mortality rate of the female heterozygous mice, we would have expected either increased seizure frequency or length as compared to males. It’s been shown that seizure burden in female mice is dependent on the phase of the reproductive cycle(29), and thus this may be contributing to the differences in seizure frequency we have reported. Others have reported differences between female and male DS mice, such as sex-dependent social interaction deficits(30) and increased mortality in females(11, 31). When examining seizure incidence and frequency, we noted that approximately 70% of seizures occur within 24 hrs of the previous seizure. Further examination of the clustering showed that 29 out of the 50 mice had evidence of clusters and on average would experience multiple seizures for 2.3 ± 1.4 consecutive days. However, on average, mice could also go approximately 4 days without seizing, with some mice having as many as 14 consecutive days between seizures (**Table 2**). Clustering adds complexity when attempting to evaluate the antiseizure efficacy of compounds. In the case where a mouse does not seize for 14 days, it may appear a coincidentally administered drug was completely protective against seizure activity, wherein this mouse may be in a seizure-free period between clusters. However, this seems to be rare, and in 50 recorded mice, this long of a duration of a seizure free period only occurred twice. In fact, the median consecutive days without a seizure occurring was 4 (**Table 2**). Additionally, the likelihood of mice having their seizure clusters and seizure-free periods synchronized would be rare and most likely will not pose a significant problem for drug screening. Thus, while one animal might be in a seizure free period, the rest of the cohort may still be seizing. Nonetheless, to capture adequate seizures for analysis and sufficiently interpret drug effects, larger sample sizes may be needed to compensate for cluster-related challenges.

Prior to approximately 76% of seizures in *Scn1a*^*A1783V/WT*^ mice, inter-ictal spiking was observed (**Figure 4**). Inter-ictal spikes are widely accepted as a diagnostic biomarker for epilepsy and may be involved in altering the balance of ion channels, allowing brain activity to cross a threshold and allow a seizure to become possible(32, 33). Spike rate prior to a seizure is also altered in humans and rodents and are related to the seizure focus(34-36). Inter-ictal spikes have been shown in another mouse model of DS to increase during hyperthermia-induced seizures(37), however if they are increased prior to a spontaneous seizure had not been examined before the present study. The increased incidence in interictal spike rates that occur shortly before a seizure should allow for prediction of impending seizures and thus, the interictal spike analysis performed here provides information for future work utilizing closed loop strategies in neuromodulation of seizures.

In a previous study, where a series of anti-seizure drugs were screened against the S*cn1a*^*tm1Kea*^ mouse model, low seizure frequency over a 60 hr recording period led to low statistical power. To overcome this limitation, an etiologically relevant priming event (a single hyperthermia-induced seizure) was added to increase spontaneous seizure frequency and subsequently increase statistical power(38). To determine if a hyperthermia-induced seizure instigated an increase in spontaneous seizure frequency in the *Scn1a*^*A1783V/WT*^ mice, we monitored spontaneous seizures over 2 weeks, administered a single hyperthermia-induced seizure, and then monitored spontaneous seizure activity for an additional two weeks. We found this single priming event did not significantly increase the average spontaneous seizure frequency in the tested cohort of *Scn1a*^*A1783V/WT*^ mice. However, two mice that had no seizures during the baseline recording period had increased seizure frequency after a hyperthermia-induced seizure. Although, clustering analysis has shown mice can go up to 14 days without seizure activity and this could be related to the clustering property we observe in *Scn1a*^*A1783V/WT*^. Thus, this priming event does not appear to be useful for significantly increasing overall seizure frequency to aid in future drug screening. However, mice underwent hyperthermia-induced seizure testing when they were well into adulthood, and at a stage when the etiological priming event may not have had as robust an effect. Additionally, as part of the protocol, heat was removed as soon as a seizure event was evident. Another study induced an acute hyperthermia-induced seizure followed by a prolonged hyperthermia-induced seizure and significantly increased the total number of seizures DS mice experienced(39). This protocol could be implemented in the future to determine if it increases the spontaneous seizure frequency. An increase in spontaneous seizure frequency could potentially reduce the number of animals required to perform a sufficiently powered study.

In patients with DS, sleep disturbances and sleep/nocturnal seizures are often reported(22, 40, 41). Similarly sleep impairment has been observed in both mouse models of DS and in scn1ab mutant zebrafish larvae(19, 42-44). Therefore, we determined the time of day when most seizures occurred in *Scn1a*^*A1783V/WT*^ mice to see if it correlated with patient reports. Interestingly, we discovered *Scn1a*^*A1783V/WT*^ have significantly more seizures between 18hr-24 hr (**Figure 6B**). The mice are housed in a room with 12-hr light/12-hr dark light cycle, with 18hr-24 hr being the first six hours during the dark cycle. Mice are nocturnal animals, sleeping primarily during the day and most active at night. Rather than most seizures occurring during sleep, as observed in humans, most seizures occurred during the *Scn1a*^*A1783V/WT*^ mouse’s perceived awake time. However, we did not analyze circadian rhythms or track sleep patterns in these mice, so we cannot say with confidence that mice were in either wake or sleep cycles when seizures occurred. Nonetheless, this provides important information about time of dosing for test compounds. Since a significant number of spontaneous seizures occur between 18hr-24hr, dosing immediately prior to this time would be beneficial in reducing the frequency of spontaneous seizures.

Clobazam, although the first line treatment of DS in human patients, has a low responder rate of ∼28%(1, 22). Given clobazam is the most commonly prescribed drug in DS, and since it both significantly decreased the temperature at which mice had hyperthermia-induced seizures in this model and significantly decreased the frequency of spontaneous seizures over a 60h time period in the *Scn1a*^*tm1Kea*^ mouse model(38), we tested various doses in the *Scn1a*^*A1783V/WT*^ mouse model. In previous work, we showed that both 5 mg/kg and 10 mg/kg single intraperitoneal (i.p.) injections of CLB significantly decreased the temperature at which *Scn1a*^*A1783V/WT*^ mice seized(26). We tested subchronic i.p. injections of 5 mg/kg CLB q.d. over 10 days and found it had no effect on the average daily frequency of spontaneous seizures. We next increased the dose to 7.5 mg/kg and again injected once daily over the course of 10 days and again found this dose had no effect on spontaneous seizures. We were surprised by these results given the effectiveness of CLB monotherapy in the *Scn1a*^*tm1Kea*^ mouse model(38, 45). However, a major difference between these studies and the one presented here was the use of drug-in-food for chronic administration, sample sizes, and length of time of experiments. In our 10-day study, the 5 mg/kg CLB group and 7.5 mg/kg CLB group, had an n= 10 and n=7, respectively, while CLB was tested over 60 h with a sample size of 16 in the *Scn1a*^*tm1Kea*^ mouse model. Post-hoc analysis showed that our study, given the small effect size, was under-powered and to see any differences, a larger sample is needed. Nonetheless, to determine if a drug reduced seizures by 80% with 80% power, using a paired analysis, a sample size of 6 would be needed. Thus, we are sufficiently powered to determine if drugs have robust effects. Drug-in-food may be a better drug administration route given the decreased stress which may arise from the daily injections and once-a-day i.p injections may not be sufficient to reach steady state therapeutic levels. A pharmacokinetic study would be useful in this regard to determine optimal dosing information. While clobazam failed as a monotherapy in our study, in the clinic, when clobazam fails, valproic acid is given in addition to clobazam. If this combination fails, then stiripentol is added on to this therapy(46). In fact, we show this add-on therapy confers significant protection against hyperthermia-induced seizures in the *Scn1a*^*A1783V/WT*^ mouse model(26). Thus, the *Scn1a*^*A1783V/WT*^ mouse will be further examined by testing similar add-on therapy approaches against spontaneous seizures. Additionally, two other drugs have recently been approved by the FDA as adjunct treatments for DS: cannabidiol (CBD) and fenfluramine (FFA)(47, 48). These will also be tested against spontaneous seizures in the future to determine their efficacy in this model of DS.

The *Scn1a*^*A1783V/WT*^ mouse model of DS recapitulates behavioral comorbidities reported in patients, excitatory and inhibitory neurons exhibit electrophysiological properties consistent with what others have reported, and excitingly, this model exhibits spontaneous seizures with a low death rate, which is beneficial for chronic drug screening. Additionally, with the information gathered here, we have found that an optimal drug dosing scheme would be to provide compounds prior to dark cycles and longer drug exposure times during chronic drug studies may be needed to overcome the complexity of seizure clustering. A full pharmacological profile against spontaneous seizures is still needed, but overall, we demonstrate face validity of the model and the potential of continuing to use the *Scn1a*^*A1783V/WT*^ mouse model for screening novel drug compounds at the contract site for the NINDS Epilepsy Therapy Screening Program.

## Methods

### Animals

Mice were grouped housed in a pathogen free facility under a 12-h light/12-h dark light cycle and had access to food and water ad-libitum. Experimental animals were generated by breeding a floxed stop male Scn1a-A1783V (B6(Cg)-Scn1atm1.1Dsf/J, Jax #026133) with a Sox2-cre (B6.Cg-Edil3^Tg(Sox2- cre)1Amc^/J) female mouse to produce both heterozygous (*Scn1a*^*A1783V/WT*^) and wildtype offspring. Both female and male heterozygous and age-matched wild type littermates were used for experiments.

### Assessing Nest Building

Nest building behavior was assessed in both male and female mice beginning at postnatal day (PND) 28. Mice were singly housed to best correlate the nest building behavior to each mouse. Prior to the onset of the experiment, mice were handled for three consecutive days for 15 minutes in the testing room to habituate them to the experimenter. Each day, for 7 days, both *Scn1a*^*A1783V/WT*^ and wild type littermates were given a 2 in^2^ cotton nestlet. Nest building abilities were scored on the rating scale as previously described(49) by an observer blinded to genotype. The score ranged from 1 to 5, where a 1 indicated the nestlet was not noticeably touched and a 5 indicated a near perfect nest. On Day 1, nesting behavior was scored at 2, 6, and 24 hours after the nestlet was added to the cage. On Days 2-7, each mouse was given a new nestlet every 24 hrs to score daily nest building.

### Open Field Paradigm

Male and female *Scn1a*^*A1783V/WT*^ and age matched wild type littermates were assessed in an open field paradigm at PND 28. Movement was automatically detected and recorded in a 60×60 cm^2^ open field arena enclosed with Plexiglass (Fusion S v1.1, AccuScan Instruments, Columbus Ohio) for 10 minutes. The behaviors scored included the total distance traveled (cm), vertical activity count, average speed (cm/s), and time spent in the center zone (s). The center zone was defined as the area 5 cm from the edge of the wall. Data was exported and analyzed in GraphPad Prism 8.0.2.

### Brain slice electrophysiology-voltage and current-clamp

Whole cell patch clamp technique was used to record from individual brain slices from both male and female *Scn1a*^*A1783V/WT*^ mice and their wild-type littermates. Slice preparation was done as previously described(50). Mice were anesthetized with pentobarbital (60 mg/kg, i.p.), brains were extracted and placed in ice-cold sucrose based ACSF continuously bubbled with 95% O_2_/5% CO_2_. Sucrose-based ACSF contained the following: Sucrose [180 mM], KCL [3 mM], Na_2_PO_4_ [1.4 mM], MgSO_4_ [3.0 mM], NaHCO_3_ [26 mM], glucose [10 mM], and CaCl_2_ [0.5 mM]. Brain slices from the dorsal hippocampus were cut with a vibrating microtome (VT1000S, Leica Microsystems Inc., Wetzlar, Germany) and transferred to an ACSF holding chamber for a minimum of 1 hour until use. For experimentation, single slices were transferred to a perfusion chamber (Warner RC-27L, Warner Instruments, Hollison, MA), anchored by a nylon net, and continuously perfused with oxygenated ACSF at room temperature. ACSF was prepared with the following: NaCl [126 mM], KCL [3 mM], Na_2_PO_4_ [1.4 mM], MgSO_4_ [1.0 mM], NaHCO_3_ [26 mM], glucose [10 mM], and CaCl_2_ [2.5 mM]. For current clamp experiments, patch electrodes were filled with potassium gluconate internal: K gluconate [132 mM], KCL [8 mM], HEPES [10 mM], EGTA (KOH) [1 mM], CaCl_2_ [0.5 mM], glucose [10 mM]. For voltage clamp recordings, patch electrodes were filled with Cs MeSO_4_ internal: Cs MeSO_4_ [140 mM], HEPES [10 mM], EGTA (CsOH) [1 mM], CaCl_2_ [0.5 mM], glucose [10 mM], ATP [2 mM], GTP [0.5 mM], QX-134 [5 mM]. Cells located in CA1 region of the hippocampus were blindly patched with borosilicate glass pipettes (2-3 MΩ). Recordings were collected using a MultiClamp 700 B amplifier and the pClamp 10 software package interfaced to an Axon Digidata 1440 A digitizer (Molecular Devices, San Jose, CA). Signals were digitized at 10 kHz and filtered at 1 kHz. To record electroresponsive membrane properties, a graded series of hyperpolarizing and depolarizing current pulses (50 pA increments, 2s in duration) were applied in current clamp mode. For recording isolated spontaneous excitatory and inhibitory post synaptic currents (sEPSCs, sIPSCs), cells were held at -70 and 0 mV, respectively; these holding potentials were used to minimize chloride-mediated and combined sodium/potassium-mediated currents, respectively. In 30 s increments, access and membrane resistance, temperature, and holding current were monitored. Cells whose access resistance exceed 40 MΩ were excluded from analysis. The amplitude and inter-event intervals for sEPSCs and sIPSCs were analyzed with Mini Analysis Program (Synaptosoft Inc., Decatur, GA, USA). 50 events recorded approximately 10 minutes after achieving whole cell configuration were randomly selected (amplitude and inter-event interval) from each cell and were averaged and compared.

### Electrode implantation surgery

Both wild type and *Scn1aA*^*1783V/WT*^ were implanted at PND 30-35 with a cortical, bipolar electrode (MS333/6-B/SPC, PlasticsOne, Roanoke, VA). Briefly, mice were anesthetized with 4% isoflurane and then transferred to a stereotaxic frame equipped with a nose cone for isoflurane delivery. Isoflurane was maintained at 2% for the duration of the surgery. One hour prior to surgery, mice were administered buprenorphine (0.01-0.2 mg/kg, i.p., MWI Animal Health, Boise, ID), and penicillin (60,000 units s.c). A hole was drilled over the skull at AP = -2.3 mm; ML = 2.55 (relative to bregma) and a bipolar electrode was placed on the surface of the dura mater, with a ground electrode placed over the cerebellum. A total of three anchoring screws were placed over the cortex, two ipsilateral of the electrode implantation sites and one contralateral. Both the electrode and the anchor screws were secured to the skull using dental acrylic (Bosworth Company, Midland, TX).

### Video-EEG Monitoring and seizure analysis

Mice recovered from the electrode implantation surgery for 14 days before being tethered for 24hr/day 7 day/week video-EEG recording. Mice (PND 45-50) were tethered to a rotating commutator (Plastics One, Roanoke, VA), which was connected to a EEG100c amplifier (BIOPAC Systems, Goleta, GA). EEG signals were digitized using a BIOPAC MP150 Recording System (BIOPAC Systems, Goleta, GA), and video was captured by DVP 7020BE Capture cards. Both the EEG data and the videos were synchronized and written to a disk using custom software(51). Video was stored in MPEG4 format, and EEG records were stored in a custom file format. EEG was reviewed by an individual blinded to genotype and drug assignment. Seizures were defined as periods of high frequency activity, 2x the baseline, lasting at least 5 seconds and followed by post-ictal depression. Average seizure frequency for each mouse was calculated by counting the number of seizures and dividing by length of time for the experiment. Individual seizure frequencies were averaged together to obtain the average seizure frequency. Seizure length was calculated as the length of the electrographic seizure in seconds. Seizure clusters were classified as previously described(21) and defined as one or more seizures per day for at least three days and at least five seizures in a cluster period. Inter-ictal spikes preceding seizures were defined as brief paroxysmal electrographic discharges at least two times the baseline values. Inter-ictal spike frequencies were calculated as the number of spikes present in the two minutes prior to a seizure.

### Hyperthermia-induced seizures

Mice were un-tethered for hyperthermia-induced seizure testing. To evaluate the temperature at which *Scn1a*^*A1783V/*WT^ mice seized, 8-week-old heterozygous mice were placed under a heat lamp and core temperature was gradually raised in a plexiglass chamber until a generalized seizure was observed or the core temperature reached 42.5°C(38). After the procedure, mice were transferred to a granite block to quickly bring the core temperature down. After approximately 5-minutes, mice were returned to their home cages. Body temperature was monitored using a neonate rectal probe (Braintree Scientific, Inc, Braintree, MA) coupled to a TCAT-2LV controller (Physitemp Instruments, Inc, Clifton, NJ). Following testing, mice were re-tethered and returned to their home cage.

### Drug Administration, i.p

*Scn1a*^*A1783V/WT*^ mice were administered clobazam (CLB) intraperitoneally (i.p). CLB (Sigma-Aldrich, Inc, St. Louis, MO) was administered in a cross-over scheme at a dose of 5 mg/kg and 7.5 mg/kg in the vehicle, 0.5% methyl cellulose. In the first leg of the cross-over, mice received either 5 mg/kg CLB or vehicle for 10 days, followed by a 7-day washout. After the washout, the treatment groups were switched and mice that previously received vehicle now received the CLB 5 mg/kg. Following another 7-day washout, CLB 7.5 mg/kg was administered in a similar manner.

### Statistical Analysis

Statistical analysis was conducted in GraphPad Prism 8.3.1 (GraphPad Software, San Diego, CA). Unpaired t-test was used for parametric data while Mann-Whitney test was used for non-parametric data. For studies in which the same mice were used for drug tests and hyperthermia-induced seizure testing, a Wilcoxon signed rank test was used. Figure legends or text contain p-values and tests used for each result. Statistical significance was defined as p < 0.05. Data is presented as mean ± SD.

### Study Approval

All animal care and experimental procedures were approved by the Institutional Animal Care and Use Committee (IACUC) of the University of Utah. Animal experiments were conducted in a manner consistent with the Animal Research: Reporting of In Vivo Experiments (ARRIVE) guidelines.

## Author Contributions

CDP, CSM, PJW, KSW designed research studies. CDP, AS, EJD, and KJJ conducted experiments and acquired data. CDP and AS analyzed data. CDP prepared the manuscript and AS, EJD, KJJ, CSM, PJW, and KSW all contributed to the editing and review of the manuscript.

## Acknowledgements

The authors thank Kyle Thomson for his assistance with video-EEG recording set-up. The authors thank the Epilepsy Therapy Screening Program at the National Institute of Neurological Disorders and Stroke for their review and comments on this manuscript. The authors would like to thank Dr. Jennifer Kearney for helpful discussions. This project has been funded in whole or in part by Federal funds from the National Institute of Neurological Disorders and Stroke, Epilepsy Therapy Screening Program, National Institutes of Health and Department of Health and Human Services, under Contract No. HHSN271201600048C (KSW), an Undergraduate Research Opportunities Scholar Grant (AS), and an American Epilepsy Society Postdoctoral Research Fellowship (CDP).

