## Supplemental Data for "Evaluation of spontaneous seizure activity, sex-dependent differences, behavioral comorbidities, and alterations in CA1 neuron firing properties in a mouse model of Dravet Syndrome"

**Table S1. Electroresponsive membrane properties of CA1 pyramidal cells (PND 18-25)**

| **Property** | **Current Step (pA)** | ***Scn1a^WT/WT^***  **(N=2; cells =4)** | ***Scn1a^A1783V/WT^***  **(N=3; cells =5)** | **p-value** |
| --- | --- | --- | --- | --- |
| **Membrane Resistance (MΩ)** | **-200** | 150.2 ± 34.2 | 158.1 ± 46.2 | 0.7847 |
|  | **-150** | 160.8 ± 30.0 | 164.8 ± 46.9 | 0.8910 |
|  | **-100** | 182.4 ± 43.5 | 182.2 ± 45.1 | 0.9934 |
|  | **-50** | 209.6 ± 53.7 | 207.4 ± 50.9 | 0.9517 |
|  | **50** | 161.7 ± 70.5 | 204.1 ± 59.8 | 0.3602 |
|  | **100** | 117.1 ± 27.8 | 153.6 ± 41.8 | 0.1792 |
|  | **150** | 110.3 ± 7.5 | 135.3 ± 30.9 | 0.1581 |
|  | **200** | 116.2 ± 30.4 | 110.2 ± 32.2 | 0.7850 |
|  | **250** | 90.0 ± 5.5 | 97.7 ± 23.9 | 0.5550 |
| **Time Constants (ms)** | **-200** | 36.4 ± 4.9 | 32.8 ± 7.9 | 0.4426 |
|  | **-150** | 38.7 ± 6.3 | 33.9 ± 5.8 | 0.2771 |
|  | **-100** | 43.1 ± 8.5 | 36.4 ± 7.4 | 0.2448 |
|  | **-50** | 45.4 ± 7.2 | 37.8 ± 6.9 | 0.1553 |
| **AP inter-event Intervals (ms)** | **50** | 30.8 ± 17.0 | 20.1 ± 16.8 | 0.4039 |
|  | **100** | 21.7 ± 5.5 | 23.8 ± 10.5 | 0.7355 |
|  | **150** | 19.2 ± 5.2 | 13.9 ± 3.6 | 0.1465 |
|  | **200** | 17.8 ± 2.5 | 11.7 ± 3.4 | 0.0283* |
|  | **250** | 15.0 ± 2.5 | 10.6 ± 2.8 | 0.0538 |

*Data presented as mean ± SD; un-paired t-test, * p<0.05*

**Table S2. Electroresponsive membrane properties of CA1 pyramdal cells (PND>30)**

| **Property** | **Current Step (pA)** | ***Scn1a^WT/WT^***  **(N=4; cells =12)** | ***Scn1a^A1783V/WT^***  **(N=4; cells =13)** | **p-value** |
| --- | --- | --- | --- | --- |
| **Membrane Resistance (MΩ)** | **-200** | 163.1 ± 48.7 | 133.6 ± 35.2 | 0.0937 |
|  | **-150** | 170.0 ± 43.6 | 145.5 ± 32.3 | 0.1223 |
|  | **-100** | 182.3 ± 50.9 | 159.9 ± 37.5 | 0.2209 |
|  | **-50** | 206.6 ± 64.1 | 176.2 ± 50.2 | 0.1985 |
|  | **50** | 210.0 ± 92.1 | 202.5 ± 57.9 | 0.8065 |
|  | **100** | 140.6 ± 54.4 | 151.3 ± 31.0 | 0.5447 |
|  | **150** | 120.0 ± 28.8 | 131.6 ± 23.3 | 0.2748 |
|  | **200** | 99.2 ± 38.7 | 109.2 ± 19.5 | 0.4171 |
|  | **250** | 93.4 ± 25.2 | 106.2 ± 20.9 | 0.1768 |
| **Time Constants (ms)** | **-200** | 31.6 ± 8.8 | 35.0 ± 4.6 | 0.2292 |
|  | **-150** | 35.4 ± 8.8 | 37.8 ± 5.7 | 0.4246 |
|  | **-100** | 39.4 ± 9.1 | 40.1 ± 6.5 | 0.8368 |
|  | **-50** | 41.9 ± 14.4 | 43.4 ± 10.4 | 0.7621 |
| **Inter-burst Intervals (ms)** | **50** | 63.9 ± 54.6 | 25.4 ± 16.1 | 0.0366* |
|  | **100** | 30.1 ± 18.4 | 20.0 ± 8.5 | 0.0967 |
|  | **150** | 25.0 ± 13.8 | 17.6 ± 5.3 | 0.0953 |
|  | **200** | 20.2 ± 8.8 | 15.0 ± 3.4 | 0.0697 |
|  | **250** | 17.4 ± 6.9 | 14.3 ± 3.7 | 0.1784 |

*Data presented as mean ± SD; un-paired t-test, * p<0.05*

**Table S3. Comparison of PND 18-25 vs. P30 *Scn1a^WT/WT^* and *Scn1a^A1783V/WT^* Electroresponsive membrane properties**

|  |  | ***P-value*** | |
| --- | --- | --- | --- |
| **Property** | **Current Step (pA)** | ***PND 18-25 vs.***  ***PND >30 Scn1a^WT/WT^*** | ***PND 18-25 vs.***  ***PND>30 Scn1a^A1783V/WT^*** |
| **Membrane Resistance (MΩ)** | **-200** | 0.5786 | 0.1652 |
|  | **-150** | 0.3905 | 0.2793 |
|  | **-100** | 0.3508 | 0.1359 |
|  | **-50** | 0.2439 | 0.1488 |
|  | **50** | 0.3543 | 0.1705 |
|  | **100** | 0.3785 | 0.3632 |
|  | **150** | 0.5872 | 0.3003 |
|  | **200** | 0.8477 | 0.4162 |
|  | **250** | 0.5243 | 0.1183 |
| **Time Constants (ms)** | **-200** | 0.2847 | 0.0702 |
|  | **-150** | 0.4272 | 0.0610 |
|  | **-100** | 0.4346 | 0.0739 |
|  | **-50** | 0.6317 | 0.0662 |
| **Inter-burst Intervals (ms)** | **50** | 0.4019 | 0.9904 |
|  | **100** | 0.6351 | 0.9789 |
|  | **150** | 0.7413 | 0.5754 |
|  | **200** | 0.9274 | 0.5637 |
|  | **250** | 0.8562 | 0.3055 |
| **Resting Membrane Potential (mV)** | **-----** | 0.3134 | 0.9553 |
| **AP Threshold (mV)** | **-----** | 0.9745 | 0.2181 |
| **Max Depolarization slope (mV/ms)** | **-----** | 0.8975 | 0.6530 |
| **Max Depolarization (mV)** | **-----** | 0.9798 | 0.3715 |
| **Max Repolarization slope (mV/ms)** | **-----** | 0.9854 | 0.5532 |

**Table S4. Sex-dependent differences in clustering**

|  | **Consecutive Days with SRS** | | **Consecutive Days without SRS** | | **# of mice with clusters/# of mice without Clusters** | | **SRS # per Cluster** | | **Cluster Duration (days)** | |
| --- | --- | --- | --- | --- | --- | --- | --- | --- | --- | --- |
| **Sex** | M | F | M | F | M | F | M | F | M | F |
| **Mean**  **± SD** | 2.4 ± 1.5 | 2.1 ± 1.2 | 4.3 ± 3.0 | 4.3 ± 3.0 | 19/8 | 10/12 | 11.2 ± 8.1 | 8.6 ± 3.2 | 3.8 ± 1.0 | 3.6 ± 0.7 |
| **Median** | 2.0 | 2.0 | 4.0 | 4.0 | ---- | ---- | 9.0 | 9.0 | 3.0 | 3.5 |
| **Max** | 6.0 | 5.0 | 14.0 | 12.0 | ---- | ---- | 35.0 | 13.0 | 6.0 | 5.0 |
| **Min** | 0.0 | 1.0 | 1.0 | 0 | ---- | ---- | 4.0 | 4.0 | 3.0 | 3.0 |
| **p-value** | 0.2726 | | 0.6657 | | ----- | | 0.3247 | | 0.6798 | |

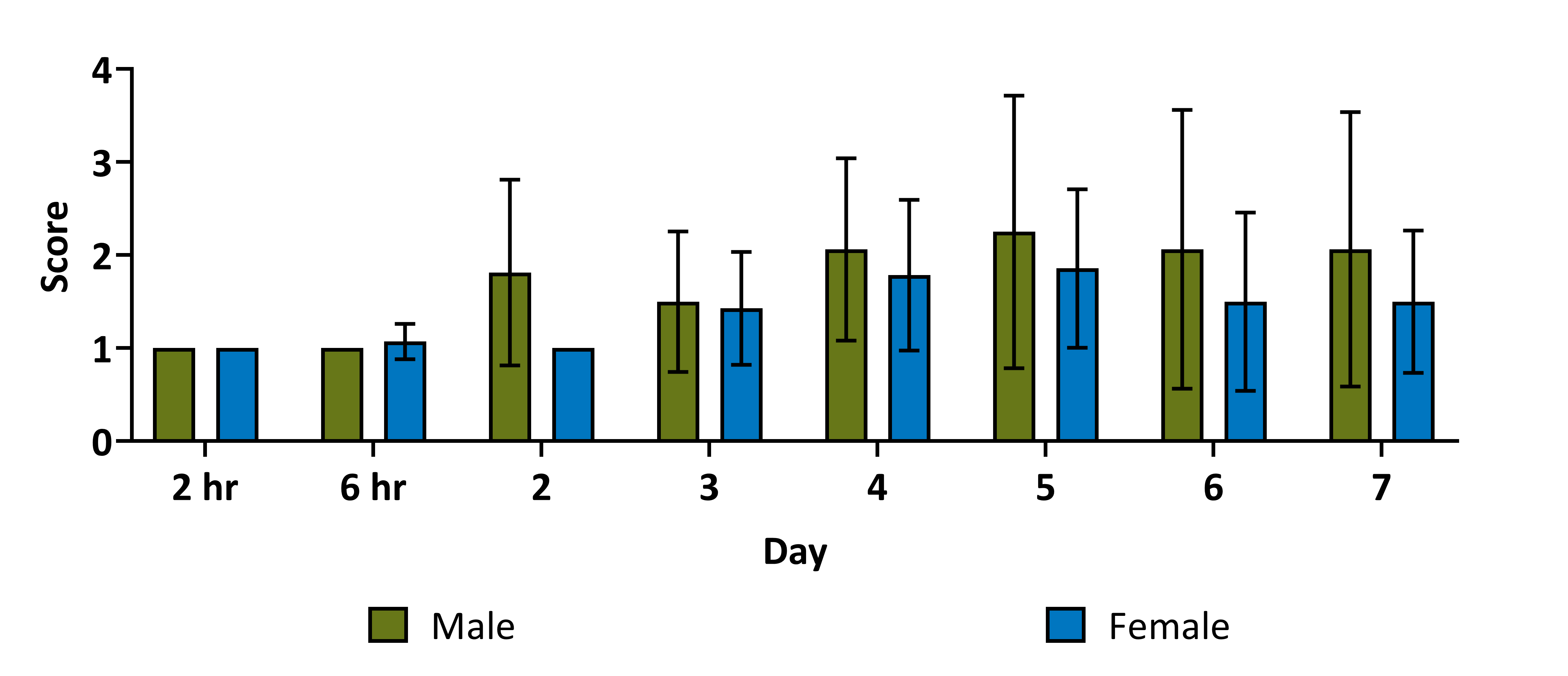

**Figure S1.** Nest building comparison between male and female PND28 –P35 Scn1a^A1783V/WT^ mice. No significant differences were present at any time-point (2 hr: p> 0.999, 6 hr: p=0.3026, Day 2: p=0.0769, Day 3: p>0.9999, Day 4: p=0.6098, Day 5: p=0.7106, Day 6: p=0.6280, Day 7: p=0.7259, Mann-Whitney, Male: n=7, Female: n=8)

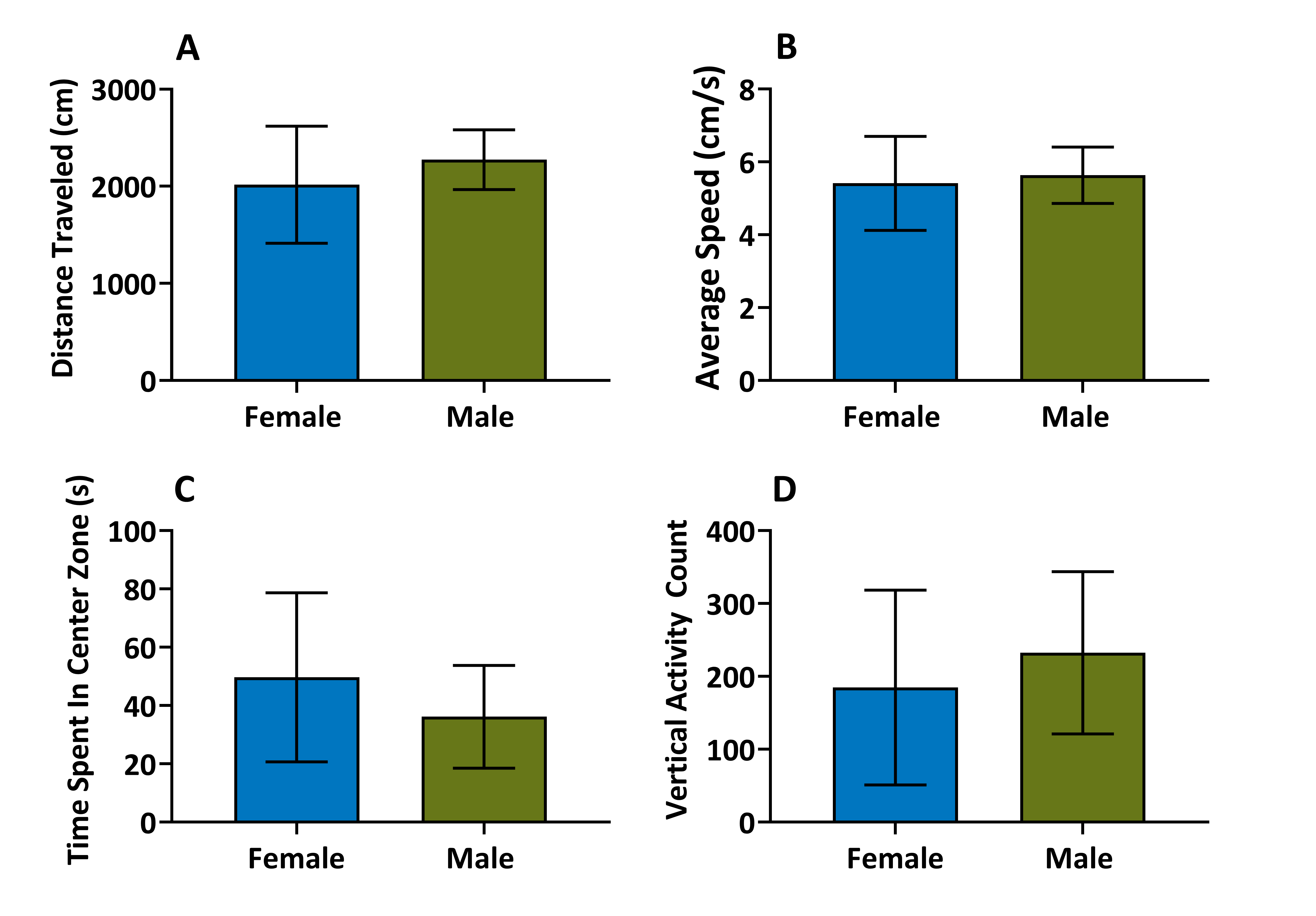

Figure S2. Open field comparison between male and female Scn1a^A1783V/WT^ mice. There were no significant differences between A) the distance traveled (p=0.3602) B) the average speed (p=0.7166), C) the time spent in the center zone (p=0.3344), or D) the vertical activity count (p=0.4923) (unpaired t-test, Male: n=8, Female: n=6)

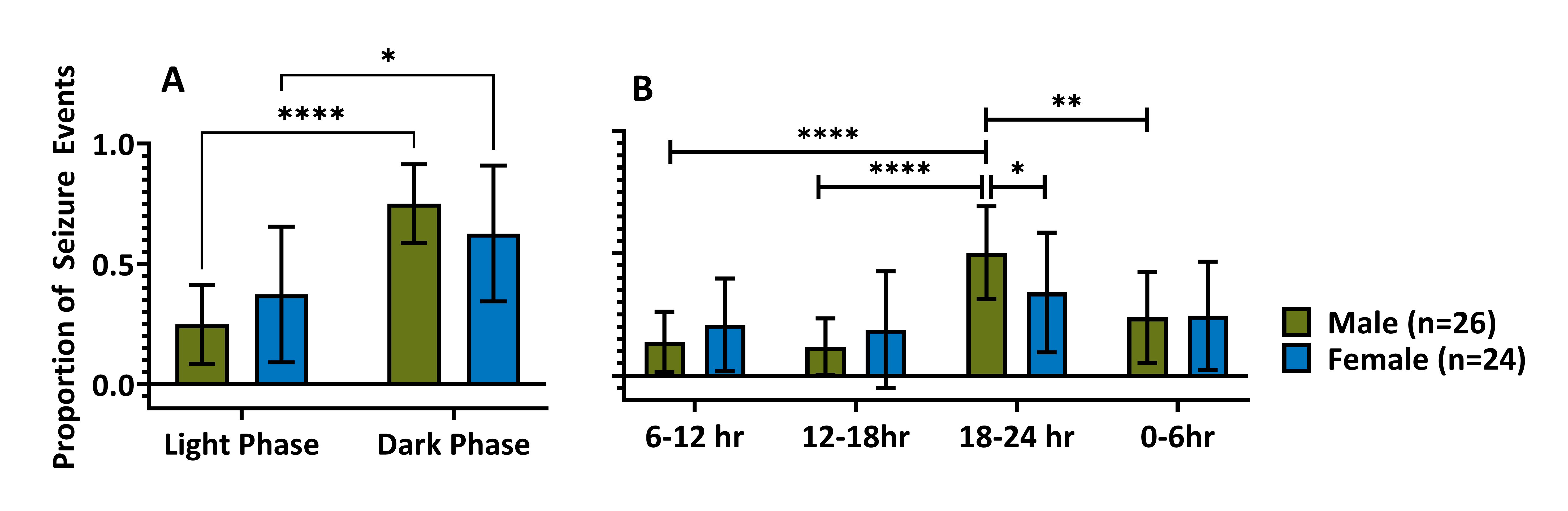

Figure S3. Time of day comparison between males and female Scn1a^A1783V/WT^ mice. A) Males and females have significantly more seizures in the dark phase as compared to the light phase (Male, p<0.0001; Female p=0.0177). B) Males seize significantly more between 18-24 hr than 6-12 hr (p<0.001), 12-18 hr (p<0.0001) and 0-6 hr (p=0.0033). There was no significant difference between the other time -points (6-12 hr vs. 12-18 hr: p=0.9953, 6-12 hr vs. 0-6 hr: p=0.2612; 12-18hr vs. 0-6 hr: p=0.1144). There was no significant difference in the time of day for females (6-12 hr vs. 12-18 hr: p=0.9999; 6-12 hr vs. 18-24 hr: p=0.3999; 6-12 hr vs. 0-6 hr: p=0.9950; 12-18 hr vs. 18-24 hr: p=0.3877; 12-18 hr vs. 0-6hr: p=0.9797; 18-24hr vs 0-6hr: p=0.7909).
